# SPEAK: Spatial Prompting with Expert Aligned Knowledge for Tissue Domain Identification in Spatial Transcriptomics

**DOI:** 10.64898/2026.06.22.733750

**Authors:** Huanhuan Wei, Xiao Luo, Hongyi Yu, Jinping Liang, Luning Yang, Lixing Lin, Maor Sauler, Naftali Kaminski, Alexandra Popa, Xiting Yan

**Affiliations:** Section of Pulmonary, Critical Care and Sleep Medicine, Yale School of Medicine, New Haven, Connecticut, USA; Yale-Boehringer Ingelheim Biomedical Data Science Fellowship Program; Department of Statistics, University of Wisconsin-Madison, Madison, Wisconsin, USA; Department of Molecular, Cell and Systems Biology, University of California Riverside, California, USA; Graduate Program in Genetics, Genomics and Bioinformatics, University of California Riverside, USA; Department of Biostatistics, Yale School of Public Health, New Haven, Connecticut, USA; Boehringer Ingelheim RCV GmbH & Co KG, Doktor-Boehringer-Gasse 5-11, 1120 Vienna, Austria; Computational Innovation, Boehringer Ingelheim Pharmaceuticals, Inc.

## Abstract

Spatially resolved transcriptomic (SRT) data requires spatial domain identification to enable tissue microenvironment-specific downstream analyses. Here we present SPEAK (Spatial Prompting with Expert-Aligned Knowledge), a large language model (LLM)-based method to identify spatial domains from SRT data by taking advantage of the prior knowledge from both LLM and human experts. SPEAK constructs a spatial context prompt for each cell/spot based on cell types and marker genes of its neighboring cells, enabling zero-shot inference, expert-guided fine-tuning, and prototype updating through two-stage prompting. Applications to STARmap, Visium, MERFISH and Xenium datasets showed advantages of SPEAK over existing spatial domain identification methods in domain prediction accuracy, robustness to limited prior knowledge, biological interpretability, and capacity for efficient expert-guided fine-tuning with generalizability to other tissue sections.

## Background

Organs and tissues exhibit complex anatomical organization, consisting of different functional units and their associated substructures. These structures are characterized by distinct cell type compositions, spatial arrangements, and molecular programs that collectively support specific physiological functions^1–4^. Consequently, understanding and dissecting these functional units is critical for identifying disease-associated alterations within a comparable structural and biological context. This will minimize confounding effects arising from differences between structures, thereby enabling more mechanistically informative insights into homeostatic processes as well as disease pathogenesis. Recent advances in spatially resolved transcriptomic (SRT) technologies^5–8^ offer powerful tools to achieve this goal. These technologies measure gene expression profiles while preserving the spatial location of cells or spatial spots within tissue sections. By simultaneously providing molecular and spatial information, SRT data provide unprecedented opportunities to identify functional units within tissues and characterize disease-associated cellular and molecular changes within each native type of functional units.

To understand the anatomical structures in SRT data, one major analytical sub-task is to identify transcriptionally homogeneous spatial domains, which are likely substructures within or even full structures of a functional tissue unit. Many methods^9–28^ have been designed for this task, which primarily use unsupervised learning algorithms on gene expression profiles, cell type annotations, and/or morphological features from co-registered histology images. A commonly used framework involves constructing a spatial neighborhood graph, generating low-dimensional cell/spot embeddings using graph neural networks, and applying clustering algorithms to partition cells/spots into distinct groups. However, these methods suffer from several limitations. First, they are unable to incorporate prior biological knowledge or expert guidance. Second, most methods conduct *de novo* training for each dataset, hindering transfer of the trained models to subjects unseen in the training dataset. Third, the unsupervised clustering results are arbitrary cluster labels lacking biological meaning, complicating interpretation and cross-sample comparisons. Finally, many methods are sensitive to the setting of the total number of clusters which is often unknown but must be specified beforehand.

To address these challenges, we developed a large language model-based method named SPEAK (Spatial Prompting with Expert-Aligned Knowledge), that leverages large language models (LLMs) to identify spatial domains from SRT data by incorporating prior biological knowledge about tissue organization from both existing scientific literatures and human experts like pathologists. We summarized the cell type compositions and gene expression profiles of the neighborhood for each cell/spot into a spatial context, which was fed into a pre-trained LLM using prompting for zero-shot spatial domain prediction. To address the limitation of pre-trained LLMs on tissues or conditions that are understudied, our method provides sample-efficient fine-tuning for the pre-trained LLM by human experts. A two-stage prompting strategy was designed to query the fine-tuned model for accurate spatial domain prediction on any given tissue section by updating domain prototypes from high-confidence predictions. We demonstrate the advantage of SPEAK through extensive applications to three benchmarking datasets with ground truth and one public real-world dataset from various tissues, species, and technologies.

## Results

### Method Overview

SPEAK is described in Methods, with the overall workflow provided in **Figure 1**. SPEAK contains three approaches: zero-shot (SPEAK-Z), fine-tuning (SPEAK-F) and two-stage prompting (SPEAK-Fs). For all three approaches, SPEAK transforms SRT data into a spatial context prompt for each cell that characterizes its neighborhood with a given size. The zero-shot approach directly feeds the spatial context prompt into a pre-trained LLM for spatial domain identification. The fine-tuning approach incorporates feedback from human experts on the zero-shot results to fine-tune the LLM for knowledge alignment. Eventually, the fine-tuned LLM agent is used to predict the final spatial domain identification results using a two-stage prompting strategy, which augmented the spatial context prompt with domain prototypes summarized from cells with high-confidence predictions. Different LLMs were used in SPEAK, including both closed-source LLMs (GPT-4o mini^29^, GPT-4o^29^, and Gemini 1.5 Pro^30^) and open-source LLMs (Llama-3.1-8B, Qwen3-30B, and Llama-3.1-70B).

**Figure 1.**
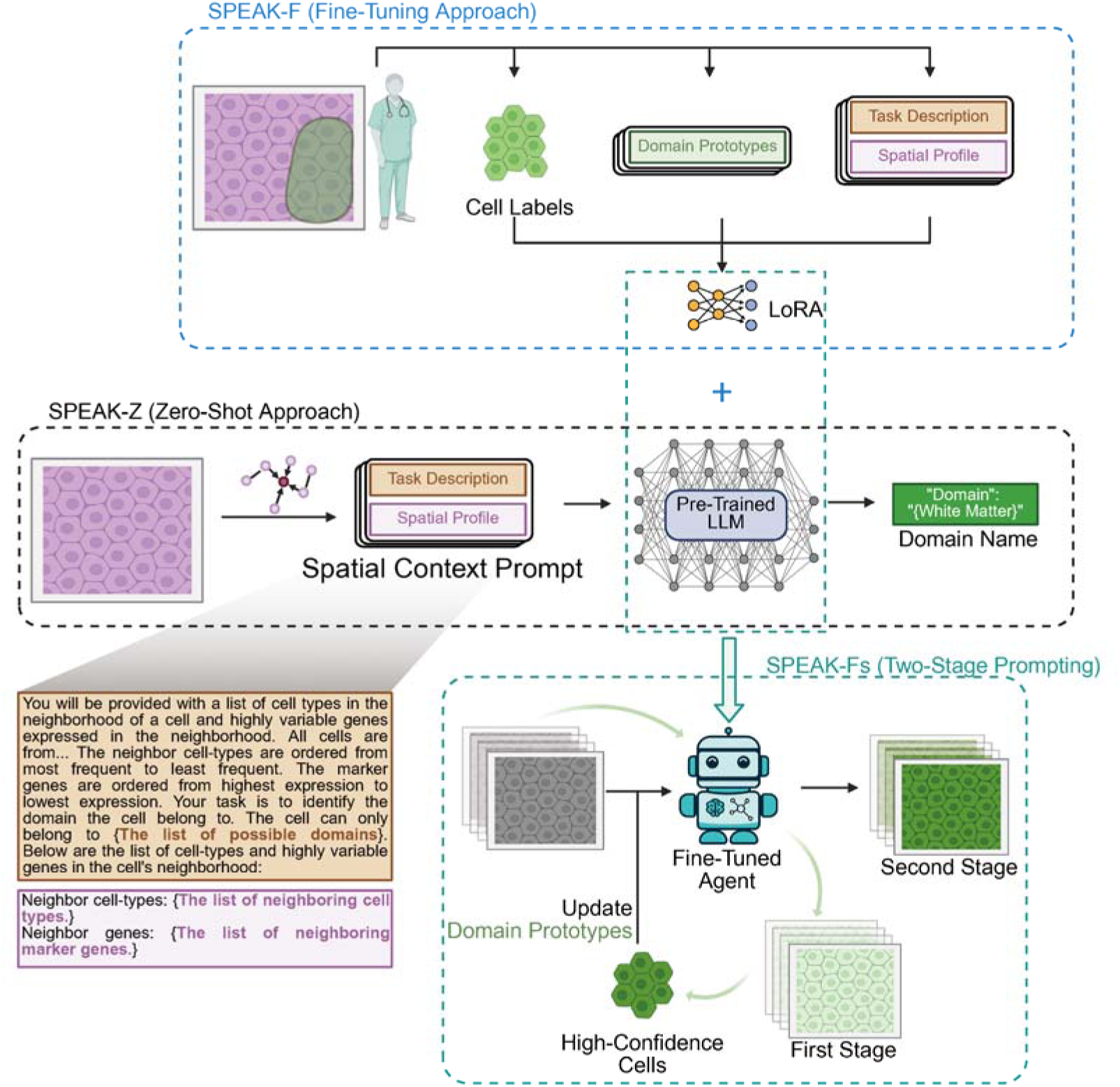
Overview of the SPEAK framework. Three key components include the zero-shot approach (SPEAK-Z), the fine-tuning approach (SPEAK-F), and the two-stage prompting strategy (SPEAK-Fs). The fine-tuning was conducted through Low-Rank Adaptation (LoRA).

### Superior Performance of Zero-shot Approach

We first benchmarked SPEAK-Z and 16 state-of-the-art spatial domain identification methods using the STARmap dataset^6^ (**Table 1**) from mouse visual cortex with single-cell resolution, containing 3 tissue sections with ground truth domain annotations for 4 different spatial domains (Layer 1, Layer 2/3, Layer 5 and Layer 6, as shown in **Figure 2c**). Comparisons between predictions of each method and ground-truth annotations using prediction accuracy-based criteria (**Figure 2a**) showed that in general SPEAK-Z had more accurate predictions than the other non-LLM based methods, with Gemini 1.5 pro and GPT-4o achieving the highest accuracy. Seurat had worse performance than SPEAK-Z but achieved the highest accuracy among the non-LLM based methods. Since Seurat conducted unsupervised clustering using cell type frequency in each cell’s neighborhood, which was the same information that SPEAK-Z used, the superior performance of SPEAK-Z over Seurat was indeed due to employing a pre-trained LLM rather than using extra or different information. In addition, we evaluated the spatial continuity of domain labels from ground truth annotation and predictions by all methods. The comparison showed that SPEAK-Z also achieved the most similar spatial continuity to ground truth (**Figure 2b**). Specifically, for both accuracy-based and continuity-based comparisons, SPEAK-Z with Gemini 1.5 Pro consistently achieved the highest performance across all criteria.

**Figure 2.**
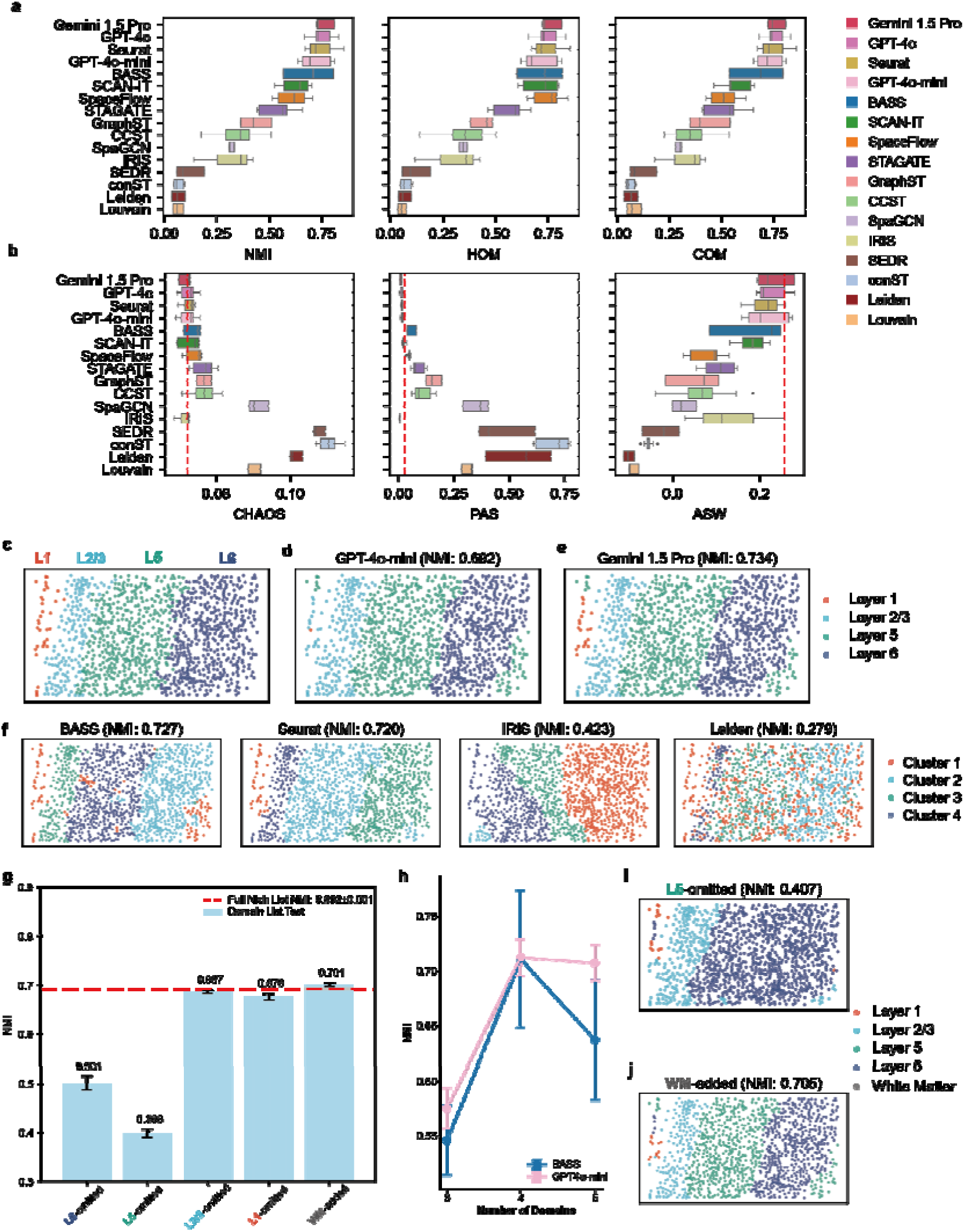
Performance evaluation and comparison of SPEAK-Z with different LLMs and other methods using the STARmap dataset. (a) Comparison using prediction accuracy-based metrics (NMI, HOM, COM) with higher values corresponding to better performance. SPEAK-Z with GPT-4o-mini, GPT-4o, or Gemini 1.5 Pro are denoted by their respective LLM names. All methods were sorted from top to bottom in decreasing order of average NMI. (b) Comparison using spatial continuity-based metrics (CHAOS, PAS, and ASW). The red dotted line indicates the spatial continuity of the ground truth. (c–f) Example tissue section showing ground truth domain (c) and domain identification results (d–f) in the STARmap dataset. (g) NMI of SPEAK-Z (GPT-4o-mini) using different potential domain lists with different types of misspecifications. (h) Comparison between SPEAK-Z (GPT-4o-mini) and BASS under different numbers of domains. (i, j) Examples of SPEAK-Z results when omitting or adding one domain.

**Table 1.** Characteristics of 4 datasets used for demonstration. The STARmap, Visium and MERFISH data have domain annotation.

|  | **STARmap** | **Visium** | **MERFISH** | **Xenium** |
| --- | --- | --- | --- | --- |
| **Tissue Type** | mouse visual<br>cortex | human dorsolateral<br>prefrontal cortex | mouse hypothalamic<br>preoptic region | human CPOD<br>lung |
| **Number of subjects** | 3 | 3 | 1 | 1 |
| **Number of replicates for**<br>**each subject** | 1 | 4 | 5 | 1 |
| **Total number of**<br>**cells/spots** | 3,268 | 47,681 | 28,317 | 9,051 |
| **Reference** | 6 | 33 | 34 | 35 |

Among the three closed-source LLMs, the performance was strongly impacted by the number of parameters in LLM (**Figure 2a-e**). Gemini 1.5 Pro has the largest number of parameters and the highest performance while GPT-4o mini has the smallest number of parameters and the lowest performance (**Supplementary Tables 1-6**). The same trend was observed among the three open-source LLMs (Llama-3.1-8B-Instruct^31^, Qwen3-30B-A3B-Instruct-2507^32^, and Llama-3.1-70B-Instruct^31^, shown in **Supplementary Figure 1**). This suggests that the performance of SPEAK may further improve as LLM technology continues to advance.

In addition to superior accuracy in grouping cells into domains, SPEAK also automatically generates meaningful labels for each identified spatial domain rather than providing abstract cluster tags with no practical meaning like non-LLM based methods (**Figure 2d-f**). This end-to-end capability saves time and labor on downstream results interpretation. Moreover, when applied to new samples, non-LLM based methods require running the entire method on the merged data containing both old and new samples, which can be computationally intensive and generate redundant but different results for the old data causing ambiguity. Instead, SPEAK can be independently applied to new samples and results from different samples can be merged directly due to the biologically meaningful domain names. Taken together, compared to the non-LLM based methods, SPEAK generates more biologically interpretable results and is computationally more efficient.

### Robustness of SPEAK to Mis-specified Domain List

One key input to SPEAK is the list of potential domains in the task description of spatial context prompt, which is partially comparable to the pre-defined number of possible domains in the non-LLM based methods such as the clustering resolution or the total number of clusters (K). Since these hyperparameters are known to significantly impact the unsupervised clustering results, we evaluated SPEAK’s robustness to the misspecifications in the domain list using the STARmap data. For computational efficiency, we used SPEAK-Z with GPT4o-mini in this evaluation. Among the non-LLM based methods, we chose BASS for comparison due to its superior performance among the non-LLM based methods (**Figure 2a,b**). We modified the ground truth domain list for SPEAK in the following ways to simulate three possible misspecification scenarios (**Supplementary Figure 2**). First, we removed one domain from the ground truth domain list for SPEAK to simulate the scenario of incomplete list. Each of the four domains was omitted separately to generate different trials so that there were only 3 domains in the list in each trial. Results across all trials were pooled together and compared to BASS for K=3. Second, we added one biologically meaningful domain (White Matter) that does not exist in the SRT data into the ground-truth domain list for SPEAK. This setting was then compared to BASS for K=5. This simulated the scenario of redundant list. Finally, we replaced one domain with “White Matter” to simulate the scenario of misnamed list.

The results showed that incomplete list reduced the performance of SPEAK but the impact diminished when the omitted domain covered a smaller area, such as Layer 1 and Layer 2/3 (**Figure 2c,g**). Similar results were observed when misnamed list was used (**Supplementary Figure 3**). However, when the redundance list was used to replace the ground-truth list (WM-added, shown in **Figure 2g**), the performance of SPEAK did not change significantly. These results emphasize the importance of including all existing domains in the list to SPEAK, regardless of whether additional redundant domains are included. Nevertheless, SPEAK was found to be more accurate and robust than BASS at the same K value (**Figure 2h**). Furthermore, omitting even a major domain that covers a large number of cells, like Layer 5, barely influenced identification of the other domains by SPEAK (**Figure 2i**). Similar results were observed when redundant list was used (**Figure 2j**). In summary, these results demonstrate the robustness of SPEAK to domain list misspecifications, which significantly reduces the need for hyper-parameter tuning by non-LLM based methods.

### Fine-tuning Approach Improves Performance

The next datasets we examined were profiled by Visium and MERFISH platforms. The Visium data (**Table 1**) measured spatial gene expression for 12 tissue sections of dorsolateral prefrontal cortex from 3 human subjects with 4 sections each at spot-level^33^. Cell type deconvolution was conducted on this data to infer cell type composition using an external reference single-cell RNA-seq dataset before SPEAK was applied. The MERFISH data (**Table 1**) measured 5 tissue sections from the hypothalamic preoptic region of a mouse at single-cell resolution^34^. The cell type annotation for each cell provided in the original study was utilized for analyses. For both datasets, we used the spatial domain annotation provided in the original study as the ground truth domain annotation. Because Gemini 1.5 Pro achieved the best zero-shot performance on the STARmap data, it was employed in zero-shot approach (SPEAK-Z) throughout the remainder of this article.

We first obtained the zero-shot results by applying SPEAK-Z to Visium data (**Figure 3a**) and MERFISH data (**Figure 3b**) separately, which demonstrated poor performance in both datasets. This is potentially due to the inaccuracy and/or low granularity of cell type annotations. The Visium data was deconvolved to estimate the cell type composition at each spot, which may be inaccurate. Besides, the cell type annotations in the MERFISH data were very broad and coarse (Excitatory and Inhibitory Neurons), providing insufficient information for domain identification. These results highlight the importance of cell type annotation quality and granularity to SPEAK-Z.

**Figure 3.**
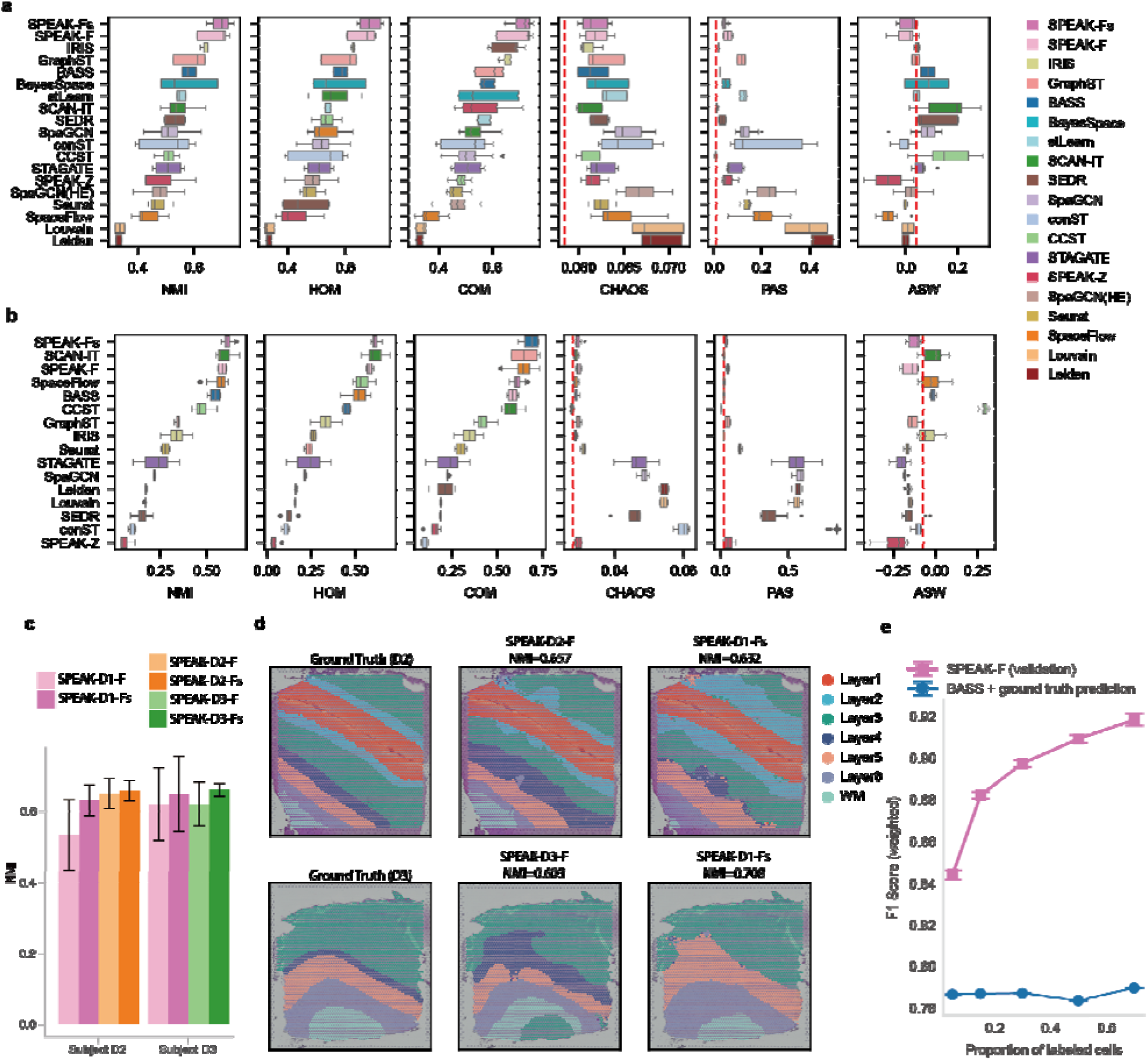
Assessment of the fine-tuning approach using Visium and MERFISH datasets. (a–b) Comparison of cluster accuracy-related metrics (NMI, HOM, COM) and spatial continuity-related metrics (CHAOS, PAS, and ASW) using the Visium (a) and MERFISH (b) datasets. The red dotted line indicates the spatial continuity of the ground truth domain labels. (c–d) Cross-subject and between-section testing results on subjects D2 and D3. (e) Comparison of prediction accuracy (F1 score) for different proportions of labeled cells revealed to SPEAK and BASS.

Due to the poor performance of SPEAK-Z, we used the Visium and MERFISH data to demonstrate the performance boost by our fine-tuning approach (SPEAK-F). For each dataset, we used 30% of cells/spots from one randomly chosen tissue section to fine-tune GPT-4o-mini, which was the only fine-tunable closed-source LLM we chose. The fine-tuned LLM (SPEAK-F) was applied to the remaining cells in the tissue section where the fine-tuning data were from (**within-section validation**) and to other tissue sections from the same subject (**between-section validation**). The between-section validation results (**Figure 3a,b**) showed that the fine-tuning significantly improved the domain predictions in both datasets, demonstrating the effectiveness of fine-tuning in addressing the limitations of the zero-shot approach when applied to spatial transcriptomic data with low cell type annotation quality. In addition, the fine-tuned model (SPEAK-F) achieved the best performance compared to all non-LLM based methods in both datasets. Similar observations were made from the within-section validation results (**Supplementary Tables 1-6**) although within-section validation had better performance than between-section validation due to variations between spatially proximal tissue sections. The within-section validation results were also robust to dropouts in gene expression (**Supplementary Figure 4**). Taken together, these results suggest that annotating a small portion of cells in a single section for fine-tuning provides sufficient information for SPEAK-F to accurately identify domains in other spatially proximal tissue sections from the same subject.

### Two-Stage Prompting Improves Cross-Subject Generalizability

To evaluate the generalizability of the fine-tuned model, we fine-tuned the model using one tissue section and applied it to a different tissue section both from the same subject (**between-section testing**) and a different subject (**cross-subject testing**). When applying the fine-tuned model, we used both the two-stage prompting strategy (SPEAK-Fs) and the regular prompting (SPEAK-F) for comparisons. For between-section testing using Visium and MERFISH datasets (**Figure 3a,b**), SPEAK-Fs had superior performance compared to all other methods, including SPEAK-F, in both datasets, suggesting the efficiency of two-stage prompting in considering differences between tissue sections from the same subject. For cross-subject testing, the Visium dataset was used which contained three subjects with four tissue slides each. We fine-tuned the model using a random tissue section from each subject (*) and applied it to all tissue sections from each other subject using two-stage prompting (SPEAK-*-Fs) and regular prompting (SPEAK-*-F). The results (**Figure 3c,d**) showed a better performance of the two-stage prompting (SPEAK-Fs) than regular prompting (SPEAK-F) again. In addition, when fine-tuning was done using a different subject (cross-subject testing), the regular prompting results on different subjects, i.e. SPEAK-D1-F on D2 and D3, had very different performance indicating strong cross-subject variance. However, the two-stage prompting results on different subjects, i.e. SPEAK-D1-Fs on D2 and D3, had similar performance. These results suggest that the two-stage prompting accommodated well for cross-subject variation and achieved more robust results. Lastly, performance of SPEAK-D1-Fs on D2 and D3 was close to that of SPEAK-D2-Fs on D2 and SPEAK-D3-Fs on D3, respectively. This shows that the two-stage prompting provides a solution for tissue sections without domain annotation for extra fine-tuning. Taken together, these results demonstrate that high-confidence cells can effectively update domain prototypes to provide tissue section-specific information without additional manual domain annotation.

### Minimal Annotation is Required for Fine-tuning

Fine-tuning required domain labeling of only a small fraction of cells to achieve high accuracy. We examined the impact of the proportion of labeled cells on fine-tuning performance using the MERFISH dataset (**Figure 3e**). Specifically, we fine-tuned SPEAK using a given proportion (5% to 70%) of cells with domain labels from a single tissue section. The weighted F1 score was calculated to measure the domain prediction accuracy for each given proportion. For the state-of-the-art (SOTA) non-LLM method, i.e. BASS, we assigned domain names to its identified cell clusters using the same percentage of labeled cells so that the weighted F1 score could be calculated (Method). In this way, both SPEAK and BASS were aware of domain labels for the same number of cells to ensure a fair comparison. **Figure 3e** shows that the accuracy of SPEAK-F surpassed that of BASS for all given proportions of labelled cells, even as low as 5%. In addition, the accuracy of SPEAK increased with increasing number of labeled cells while the accuracy of BASS stayed unchanged because it could not learn from additional labeled data. This means that SPEAK can adopt increasingly available domain annotation for a continuously increasing performance. Finally, the slope of performance gain slows down significantly when the proportion of labeled cells reached 30%. This suggests that labeling a relatively small portion of cells is sufficient to adapt a general LLM into a specialized model with high accuracy.

### Human-in-the-loop Successfully Corrects Spatial Domain Predictions in Human COPD Lung

We applied SPEAK to a Xenium dataset^35^ from the lung of a chronic obstructive pulmonary disease (COPD) patient, which offered single-cell resolution and higher gene detection capabilities and was generated using the newest generation of SRT technologies. For comparison, we also applied two existing domain identification methods, Seurat^36^ (BuildNicheAssay function) and IRIS^18^, to this dataset. All three methods had identical input: cell type annotations and spatial locations. Based on brief pathological analysis of the tissue section, potential domains in this COPD lung tissue section included eight types: Airway Inflamed, Alveolar, Fibrotic, Immune, Inflamed Other, Large Vessel Healthy, Large Vessel Inflamed, and Repair. For SPEAK, we included all 8 domain names in the spatial context prompt. For Seurat and IRIS, we set the number of clusters to 8.

Comparing results from all three methods (**Figure 4a-c**), we found two regions that were predicted to be inflamed airways (in pink square) and large inflamed vessel (in green square) by SPEAK-Z. Both Seurat and IRIS identified them to be the same domain type. Examination of cell type annotation in these two regions revealed that both regions were enriched in smooth muscle cells (SMCs), which was confirmed by the expression of ACTA2, a marker of SMCs (**Figure 4d**). However, region in pink square reside in a larger structure that is enriched for expression of FOXJ1 (a ciliated cell marker) and TP63 (a basal cell marker), which are major cell types in airways instead of in vessels. This indicates that the regions in pink and green squares are smooth muscle cells from airway and vessel, respectively. Histological analyses of the H&E staining image (**Figure 4a**) confirmed these results. These results showed that SPEAK is more sensitive to subtle differences in the neighborhood of cells. Potentially, changing the number of clusters in Seurat and IRIS can achieve similar results but this parameter tuning is impossible for tissue sections without any prior domain annotation knowledge.

**Figure 4.**
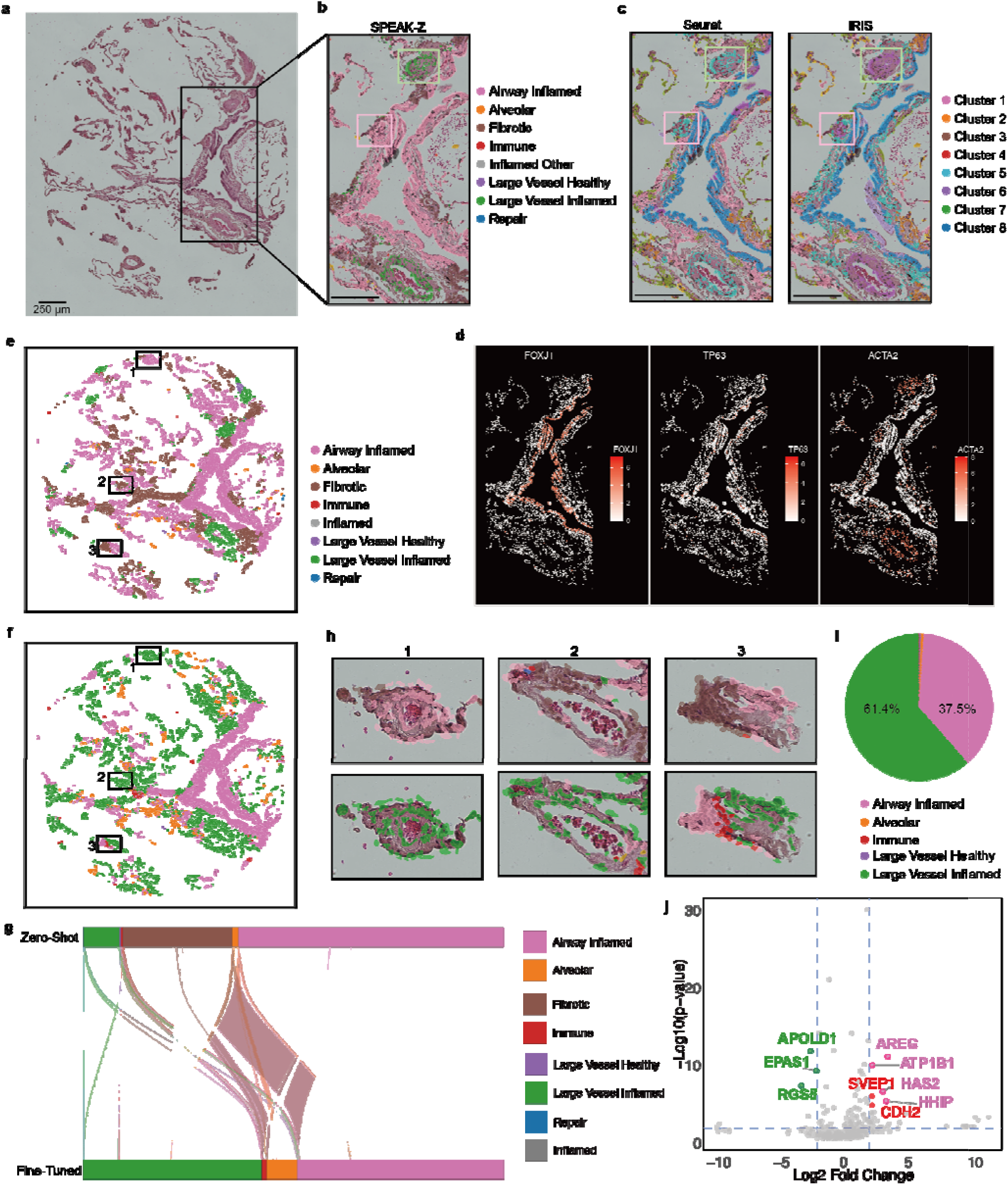
Application of SPEAK to a Xenium human COPD lung dataset. (a) H&E image of the human COPD lung sample. (b–c) Domain identification results in a selected region comparing SPEAK-Z to Seurat and IRIS. (d) Expression levels of marker genes in the selected region. (e) Domain identification results by SPEAK-Z. (f) Domain identification results by SPEAK-F. (g) Alluvial diagram of domain identification results comparing SPEAK-Z and SPEAK-F. (h) Three selected regions illustrating differences between SPEAK-Z and SPEAK-F. (i) Pie chart showing the proportion of smooth muscle cells (SMC) assigned to different domains by SPEAK-F. (j) Volcano plot of differentially expressed genes between SMCs in the Airway Inflamed and in the Large Vessel Inflamed. Genes with absolute log2 fold change larger than 2 and adjusted p-value < 0.05 are colored. Canonical airway-related and highlighted in red.

In addition to the zero-shot approach (SPEAK-Z), we fine-tuned the model (SPEAK-F) using cells from the region in **Figure 4b** with consistent annotation from SPEAK-Z and histological analyses. Comparison between fine-tuning (**Figure 4f**) and zero-shot (**Figure 4e**) results showed that most cells from Large Vessel Inflamed and Airway Inflamed remained in the same category after fine-tuning (**Figure 4g**). Meanwhile, many cells identified as Airway Inflamed or Fibrotic by SPEAK-Z were re-labeled as Large Vessel Inflamed (**Figure 4g**) by SPEAK-F. To determine whether the SPEAK-F results were more accurate, we selected several regions with different results between SPEAK-Z and SPEAK-F and examined their H&E staining image-based histological annotation. Three examples of these regions were shown in **Figure 4h**. The H&E staining image-based histological analyses confirmed that SPEAK-F results were correct for all regions. In particular, for the region 3 in **Figure 4h**, SPEAK-Z identified the left part of the tissue as Fibrotic and the right part as Airway Inflamed. SPEAK-F successfully corrected the left part to Airway Inflamed and the right part to Large Vessel Inflamed, confirmed by the H&E staining image.

We further stratified SMCs into two groups (**Figure 4i**) based on their predicted domains by SPEAK-F: Airway Inflamed and Large Vessel Inflamed. Differential expression analysis between the two groups revealed that SMCs from the Large Vessel Inflamed domain had increased expression of canonical markers of vascular tone (RGS5) and hypoxia response (APOLD1, EPAS1) (**Figure 4j**). Instead, SMCs from the Airway Inflamed domain displayed significantly higher expression of genes essential for airway morphogenesis (HHIP)^37^, epithelial repairing (AREG), and contractility regulation (HAS2, ATP1B1). These results further confirm the accuracy of SPEAK-F predictions. Furthermore, SPEAK captured high expression of the adhesion molecule CDH2 and the ECM protein SVEP1 in SMCs from the airway domain (**Figure 4j**). While these markers are typically associated with vasculature in healthy tissue, their upregulation in airway smooth muscle cells is consistent with pathological airway remodeling and mesenchymal activation characteristic of COPD^38^. These findings demonstrate that SPEAK not only correctly distinguishes anatomical domains but also identifies context-dependent pathological changes.

### Computational Efficiency and Interpretability

SPEAK is computationally efficient because spatial context prompts for all cells can be processed in parallel. When using open-source LLMs with vLLM on 2 NVIDIA H200 GPUs and 2 CPU cores, SPEAK speed ranged from 3 seconds per 1,024 cells (Qwen3-30B-A3B-Instruct-2507) to 18 seconds per 1,024 cells (Llama-3.1-70B-Instruct). When using the OpenAI API, processing time depended on network conditions and service load. In general, obtaining results for a STARmap sample with 1,088 cells took 5 to 10 minutes. In addition, due to parallel processing, SPEAK can avoid out-of-memory errors. Some non-LLM methods, especially the graph-based ones, can be infeasible due to memory limitations as cell counts increase, whereas SPEAK remains applicable as long as the LLM can be loaded on the GPU. Lastly, SPEAK has the ability to output reasoning steps for domain predictions (**Supplementary Figure 5**). This makes it more interpretable than other non-LLM methods. The reasoning steps can help biologists evaluate whether the LLM is applying correct knowledge, providing both a diagnostic tool and mechanism understanding when applying SPEAK to real data.

## Discussion

We have developed a novel human-in-the-loop spatial domain identification method, SPEAK, that takes advantage of pre-trained large language models with fine-tuning to identify spatial domains from spatial transcriptomic data. Through applications to 3 public benchmarking SRT datasets and 1 validating dataset that were from various SRT technologies, species, tissues and conditions, we demonstrated the superior performance of SPEAK to current non-LLM spatial domain identification methods across multiple accuracy and continuity metrics under the zero-shot setting. The fine-tuning and two-stage prompting strategy in SPEAK further improved its performance and achieved strong generalizability across tissue sections and subjects. The method combines accuracy, interpretability, and practical efficiency, offering a scalable solution where minimal human annotation yields substantial results across diverse spatial transcriptomic platforms.

SPEAK offers several practical advantages over existing methods. First, it takes advantage of the biological knowledge encoded in both pre-trained LLMs and human experts, and generate biologically meaningful domain names rather than abstract cluster indices. This eliminates the need for post-hoc annotation and enables direct comparison across batches without re-running clustering procedures. When cell type names were replaced with abstract labels without biological meaning, the performance of SPEAK-Z dropped substantially (NMI from 0.72 to 0.19), confirming that LLMs leverage semantic understanding of biological terminology rather than pattern matching alone. Second, SPEAK performance will continue to improve as LLM technology advances. Our results showed that larger models consistently achieved better performance in zero-shot tasks in both proprietary (GPT-4o mini vs. GPT-4o) and open-source models (Llama-3.1-8B vs. Llama-3.1-70B). Third, SPEAK offers a highly scalable solution where minimal human guidance can yield substantially accurate results with excellent generalizability. The human guidance can also put more emphasis on wrongly predicted domains by selectively providing more fine-tuning data to maximize the efficiency of fine-tuning. Fourth, different from non-LLM methods, SPEAK tolerates both missing and redundant domain names in the potential domain list, making it applicable to less-characterized tissues where the exact number of domains is unknown. Fifth, SPEAK is computationally efficient due to its parallel processing of individual cells to avoid the memory limitations which affects graph-based methods as cell counts increase. Sixth, SPEAK can output the reasoning steps to help biologists evaluate whether the LLM utilizes the correct knowledge, providing both diagnostic and mechanism understanding tools. This interpretability distinguishes SPEAK from other deep learning methods that embed cells into abstract latent spaces.

SPEAK has a few limitations that need future work. First, it strongly relies on the previous knowledge encoded in the large language model. For tissues and conditions that are understudied, the performance can be limited. Potentially, confidence score of predictions by LLM can be used to identify the under studied cases which would require more human guidance. Second, a list of potential domains existing in the data is needed and important to SPEAK, requiring prior knowledge about the tissue section. Future work can assist in generating this list directly from pre-trained LLMs. Third, the size of cell neighborhood is currently pre-defined in SPEAK, which may have different optimal settings for different tissues, domains, or cell types. Future work can focus on automatically identifying the optimal neighborhood size. Lastly, our results showed that larger LLMs are more challenging to fine-tune. Future development can pursue knowledge distillation to smaller, open-source architectures, enabling deployment in resource-constrained environments.

## Methods

### Overview of SPEAK

The overall workflow of SPEAK can be found in Figure 1, which contains three approaches: zero-shot (SPEAK-Z), fine-tuning (SPEAK-F) and two-stage prompting. For all three approaches, SPEAK transforms spatial transcriptomic data into spatial context prompt consisting of a spatial profile and task description for each cell. The spatial profile is constructed using the cell type composition and average gene expression profiles of neighboring cells located within a pre-defined distance from each given cell. For SRT data with spot resolution, we consider each spot as one cell with its cell type proportion as the cell type annotation so the same procedure can be applied to construct the spatial profile. The task description provides the list of possible target domains to identify, with an optional response format instructions to enable output reasoning details.

In the zero-shot approach, the spatial context prompt for each cell was directly fed to a pre-trained LLM for spatial domain identification. Based on the zero-shot results, the fine-tuning approach provides an interface for human experts to guide the LLM for improved predictions by providing feedback on selected cells with inaccurate prediction. Specifically, the correct domain labels for chosen cells are provided by human experts, based on which domain prototypes are generated by averaging the spatial profiles of labeled cells from the same domain. The domain labels and spatial context prompts of the chosen cells, together with the generated domain prototypes, are used for low-rank adaptation fine tuning (LoRA^39^). Eventually, the fine-tuned LLM agent will be used to generate the final spatial domain identification results using a two-stage prompting strategy. In the first stage, the fine-tuned agent generates initial domain predictions for unlabeled cells, based on which the sample-specific domain prototypes are calculated using cells with high-confidence predictions. In the second stage, the fine-tuned agent will use the sample-specific domain prototypes, and the spatial context prompts to generate the final spatial domain identification.

### Spatial Context Prompt Construction

Suppose the SRT dataset 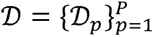 include data from *P* subjects with 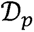 denotes the SRT data for subject *p*. For subject *p*, let 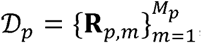, where **R***_p,m_* denotes data of the tissue section *m* and *M_p_* represents the total number of tissue sections from subject *p*, respectively. **R***_p,m_* = (**X***_p,m_*, **S***_p,m_*, **c***_p,m_*) consists of the following 3 components:

1. Spatial Coordinates: 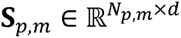, where *d* is the dimensionality of the spatial coordinates (typically *d* = 2) and *N_p,m_* is the total number of cells/spots on the section.
2. Cell Type Annotations: 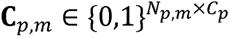, where *C_p_* is the total number of cell types in subject *p*, row *i* of ***C****_p,m_* is a one-hot encoded vector representing the cell type annotation for cell/spot *i*. Note that, for spot-level SRT data like the Visium dataset, each row is a vector of the cell type proportion from the deconvolution results.
3. Cell Type Marker Gene Expression Matrix: 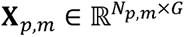, where *N_p,m_* is the number of cells and *G* represents the total number of cell type marker genes, which were selected in the following way. For SRT data with single-cell resolution (MERFISH, STARmap, and Xenium), data from different sections were integrated using canonical correlation analysis (CCA) and clustered using Leiden clustering (resolution=0.1). The top five positive marker genes for the identified clusters were selected as the cell type marker genes. For SRT data with spot-level resolution (Visium), we use established cell type marker genes from existing literatures.

A spatial context prompt is constructed for each cell/spot *i* comprised of 2 components including the task description and spatial profile (**Figure 1**). The task description specifies the objective, species, tissue type, names of potential spatial domains existing in the data and format of input and response. By default, our response format requires the LLM to output only the most probable spatial domain without explanation. For improved interpretability, users can request detailed reasoning steps by removing the response format.

The spatial profile of cell/spot *i* is constructed to summarize the information of its neighborhood 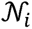 which is the set of cells located within a circle with a radius of *δ* and cell/spot *i* as its center. *δ* is set to 72 µm, 100 µm, 344 µm and 50µm for STARmap, MERFISH, Visium and Xenium data, respectively. These thresholds are selected such that the resulting neighborhood represents the minimal functional unit of the tissue domains, encompassing the characteristic scale of its structural complexity. For each neighborhood 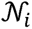, the spatial profile contains two ordered lists as follows.

a. Neighbor Cell Type List: list of cell type names sorted in descending order of their cell type frequency in neighborhood 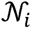, where cell type frequency is the proportion of cells belonging to each given cell type.
b. Neighbor Marker Gene List: list of marker gene names sorted in descending order of their average expression level across all cells/spots in neighborhood 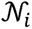.

### Zero-shot approach (SPEAK-Z)

For the zero-shot approach (SPEAK-Z), we directly fed the spatial context prompt of each cell to a pre-trained LLM. We evaluated two categories of LLMs. The first category includes those accessible through official APIs, including GPT-4o mini, GPT-4o, and Gemini 1.5 Pro. Default generation hyperparameter setting was used. The second category includes the open-source LLMs that can be run locally to ensure data privacy and sovereignty, including Llama-3.1-8B, Qwen3-30B, and Llama-3.1-70B. The pre-trained models were downloaded from HuggingFace and deployed locally using vLLM (v0.8.5).

### Fine-tuning approach (SPEAK-F)

For the fine-tuning approach (SPEAK-F), we obtain a subset of cells with spatial domain labels as the fine-tuning data. For each labelled cell in this data, SPEAK-F constructs an augmented spatial context prompt with domain prototypes defined as the average spatial profiles across all cells within each domain (**Figure 1, Figure S1**). Specifically, the cell type frequencies and average marker gene expression levels for the neighborhood of all cells within each annotated spatial domain were further averaged, based on which a list of cell type names and a list of marker gene names were constructed like how the spatial profile was constructed for each cell neighborhood. The augmented prompts together with the cell domain labels are used to fine-tune the LLM. We applied LoRA^39^ via OpenAI’s fine-tuning API for the fine-tuning which was trained for 3 epochs with a batch size of 2 and a default learning rate.

In benchmarking datasets (STARmap, MERFISH and Visium), we randomly selected 30% of cells/spots for fine-tuning. In the demonstration dataset (Xenium), we collaborated with pathologists to choose 1,187 out of 9,051 cells with correct predictions by the zero-shot approach for fine-tuning.

### Two-stage prompting (SPEAK-Fs)

To apply the fine-tuned model to other samples, we proposed a two-stage prompting strategy (SPEAK-Fs) to account for differences across samples. In the first stage, a fine-tuned model generates initial predictions. Cells with more than 50% of neighbors sharing the same predicted domain are classified as cell with high-confidence predictions. These cells are used to update the domain prototypes. For domains with no cells with high confidence predictions, we will retain the domain prototypes from SPEAK-F. In the second stage, the updated domain prototypes together with the spatial context prompts are fed into the fine-tuned model to generate improved predictions for the remaining cells without high-confidence predictions.

### Post-prediction refinement

To enforce spatial smoothness of the spatial domain predictions, we applied a post-prediction refinement step similar to SpaGCN and GraphST. We compare the prediction of each cell with that of its neighbors defined in the spatial profile construction. If more than half of its neighbors were predicted as a different domain, the cell will be relabeled to the most popularly predicted domain across its neighbors. For the zero-shot approach, we applied this step after obtaining LLM predictions. For the fine-tuned model, this step was applied after SPEAK-F or SPEAK-Fs.

### Method performance evaluation and comparison

We benchmarked SPEAK against 14 state-of-the-art spatial clustering methods and 2 non-spatial clustering methods. Each method was run three times per tissue section per subject to evaluate performance variation. The spatial clustering methods included graph-based approaches (CCST^28^, GraphST^15^, STAGATE^40^, SpaGCN^10^, SEDR^27^), integrative methods (Seurat^36^, IRIS^18^, BASS^41^, conST^25^, SpaceFlow^24^, SCAN-IT^42^, stLearn^26^, BayesSpace^11^), and a histology-augmented variant SpaGCN(HE)^10^. The non-spatial clustering methods were Leiden and Louvain. BayesSpace, stLearn, and SpaGCN(HE) require histological images and thus could not be evaluated on the STARmap and MERFISH datasets. The parameters for these baseline methods were optimized in prior benchmarking work^43^, and we directly used the reported optimal results for comparison.

The method performance was evaluated by comparing results to ground truth in the benchmarking datasets (STARmap, MERFISH and Visium) using three metrics: Normalized Mutual Information (NMI), Homogeneity score (HOM), and Completeness score (COM). All three metrics range from 0 to 1, with higher values indicating better agreement between predicted and true cluster assignments. These metrics were calculated using the scikit-learn^44^ python package. In addition to the similarity with ground truth, the spatial continuity of predictions was also assessed for method performance comparison. Three metrics were used including spatial chaos score (CHAOS^45^), the Percentage of Allowed outliers for Segmentation (PAS^45^), and the Average Silhouette Width (ASW^46^). Lower CHAOS and PAS values indicate higher spatial continuity. ASW is rescaled to 0–1, with higher values corresponding to greater spatial coherence. These metrics were calculated using SDMBench^43^.

To evaluate how the proportion of labelled cells affected fine-tuning performance, we performed an additional MERFISH analysis using labelled subsets from a single tissue section. Because BASS is an unsupervised spatial clustering method and does not directly output domain labels, we added a label-assignment step before calculating weighted F1 scores for its comparison with SPEAK-F. In the MERFISH dataset, SPEAK was fine-tuned using labelled subsets from a single tissue section, with labelled-cell proportions ranging from 5% to 70%, and weighted F1 scores were calculated by comparing its domain predictions with ground-truth labels. For BASS, we kept its clustering result unchanged and used only the same revealed labelled cells to map clusters to domains. Each BASS cluster was assigned the domain label represented by the majority of labelled cells within that cluster, after which all cells in the cluster inherited this assigned domain label for weighted F1 calculation. This protocol allowed BASS and SPEAK-F to use domain labels from the same number of cells while preserving BASS as a clustering-only baseline.

### Noise Resilience Experiments

To evaluate the robustness of SPEAK to dropouts, we randomly chose a given proportion of gene expression levels in MERFISH data and set them to 0. The proportion of randomly chosen intensities vary from 10% to 80%. For each given proportion, the top-10 neighbor marker gene list in the spatial profile constructed before and after introducing dropouts for each cell are compared to calculate the number of non-overlapping genes. This number was averaged across all cells to compute the Mean Changes in Top-10 Genes, for which higher values indicate larger impact of dropouts on SPEAK. In addition, we applied the fine-tuned model to cells not included in the fine-tuning data before and after introducing dropouts. The relative NMI drop, defined as 1 − (NMI after noise / NMI before noise), was calculated as the performance loss due to dropouts.

### Real Data Analysis

We applied SPEAK to four spatial transcriptomic datasets that span a range of technologies, species and tissue types (Table 1). When multiple tissue sections were collected from the same subject, they were taken from spatially proximal locations. The STARmap, Visium and MERFISH datasets all carry manual spatial domain annotations as the ground truth for benchmarking. The Xenium dataset of chronic obstructive pulmonary disease (COPD) lung served as a demonstration.

#### STARmap dataset

The STARmap dataset^6^ profiles the mouse visual cortex at single-cell resolution. It comprises three subjects with one tissue section each, profiling a total of 3,268 cells. The cell type annotations and the ground-truth domain annotation for the 4 cortical-layer domains (Layer 1, Layer 2/3, Layer 5 and Layer 6) were taken from the original publication. The data were preprocessed following descriptions in the original publication as well. Cell type marker genes were identified as described in the subsection of “Spatial Context Prompt Construction”. For each cell, the spatial profile excluded the marker genes and used only its neighborhood cell-type composition.

#### Visium dataset

The Visium dataset^33^ profiles the human dorsolateral prefrontal cortex (DLPFC) at spot resolution with a spot size of ∼55 µm. It comprises three subjects with four tissue sections each, providing 12 tissue sections and 47,681 spots in total. The original publication provides ground-truth annotations for seven spatial domains, including the six cortical layers and the white matter (Layer 1, Layer 2, Layer 3, Layer 4, Layer 5, Layer 6 and WM). The Visium platform does not provide cell-type annotations, so each spot was deconvolved using SDePER^47^ to estimate its cell-type composition with an independent single-nucleus RNA-seq reference data from the human DLPFC^48^ as the reference data. The estimated cell type composition of each spot was taken as its cell type annotation so that the cell type frequency of a neighborhood was calculated as the average cell type composition across the neighboring spots. In addition, 22 established DLPFC marker genes from the literature^43^ were used to construct the neighbor marker gene list in the spatial context prompt.

#### MERFISH data

The MERFISH dataset^34^ profiles the mouse hypothalamic preoptic region at single-cell resolution. It comprises one subject with five tissue sections, profiling 28,317 cells in total. The cell type annotations and the ground-truth domain annotation for eight domains (MPA, PVH, BST, PVT, MPN, PV, V3 and fx) were taken from the original publication. Since the provided cell-type annotations are coarse, neighbor marker-gene expression was added to the spatial context prompt, with marker genes selected as described in the subsection of “Spatial Context Prompt Construction”.

#### Xenium data

The Xenium dataset profiles human COPD lung at single-cell resolution^35^. We validated our method on a tissue section of 9,051 cells. No pre-existing spatial-domain annotation is available, so this dataset served as a demonstration. The eight candidate domains supplied to SPEAK were defined from pathological assessment of the tissue (Airway Inflamed, Alveolar, Fibrotic, Immune, Inflamed Other, Large Vessel Healthy, Large Vessel Inflamed and Repair). The cell-type annotations were provided by the original study. The data were preprocessed following that study. Only the cell-type composition was used to construct spatial profiles for each cell, as the Xenium panel only captures 480 genes.

## Availability of data and materials

### Data availability

The benchmarking datasets (STARmap, Visium and MERFISH) and their ground-truth spatial-domain annotations are available from the SDMBench repository on figshare (https://figshare.com/projects/SDMBench/163942). The Xenium COPD lung dataset is available from the Gene Expression Omnibus under accession GSE313006.

### Code availability

SPEAK is freely available under the MIT license at GitHub (https://github.com/wJDKnight/SPEAK). The same repository contains the processed data and the scripts needed to reproduce all analyses presented in this study.

## Acknowledgements

We gratefully acknowledge the Yale–Boehringer Ingelheim Biomedical Data Science Fellowship Program for its support. H. W. is supported by the Yale–Boehringer Ingelheim Biomedical Data Science Fellowship.

## Contributions

X.Y. conceived the idea and provided funding support. H.W., X.L. and X.Y. designed the study. H.W., H.Y., J.L., L.Y., and L.L. developed the method, implemented the software and performed simulations. H.W., A.P. and M.S. conducted the real data analysis. M.S. and N.K. provided the COPD dataset and aided result interpretation. H.W. and X.Y. wrote the manuscript. X.Y. supervised the research. All authors read and approved the final manuscript.

