## Supplemental Materials for "SPEAK: Spatial Prompting with Expert Aligned Knowledge for Tissue Domain Identification in Spatial Transcriptomics"

[*]Corresponding author

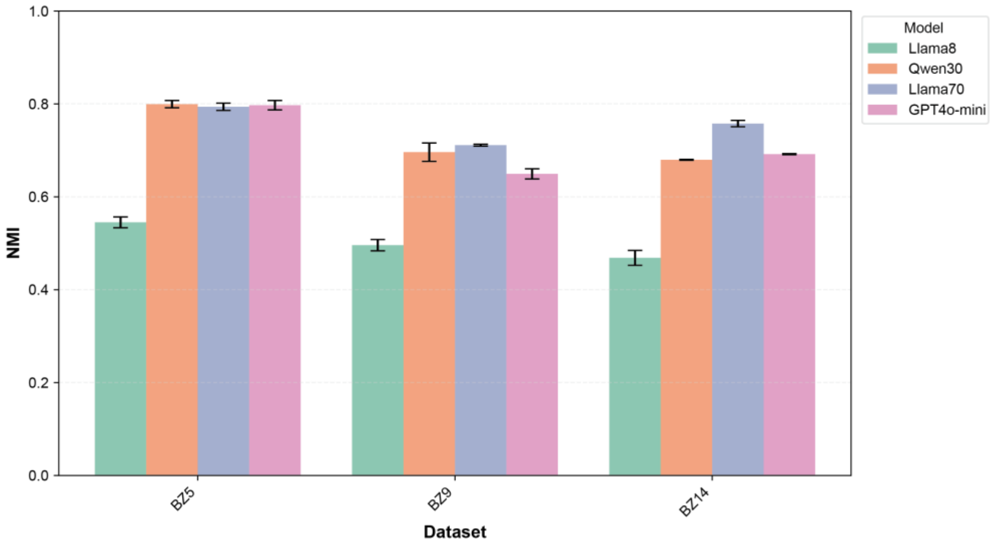


**Supplementary Figure 1.** SPEAK performance across different LLM backbones. NMI scores of SPEAK when applied to BZ5, BZ9 and BZ14 using Llama-3.1-8B-Instruct, Qwen3-30B-A3B-Instruct-2507, Llama-3.1-70B-Instruct and GPT-4o-mini are shown in bars with error bars indicating the standard error.

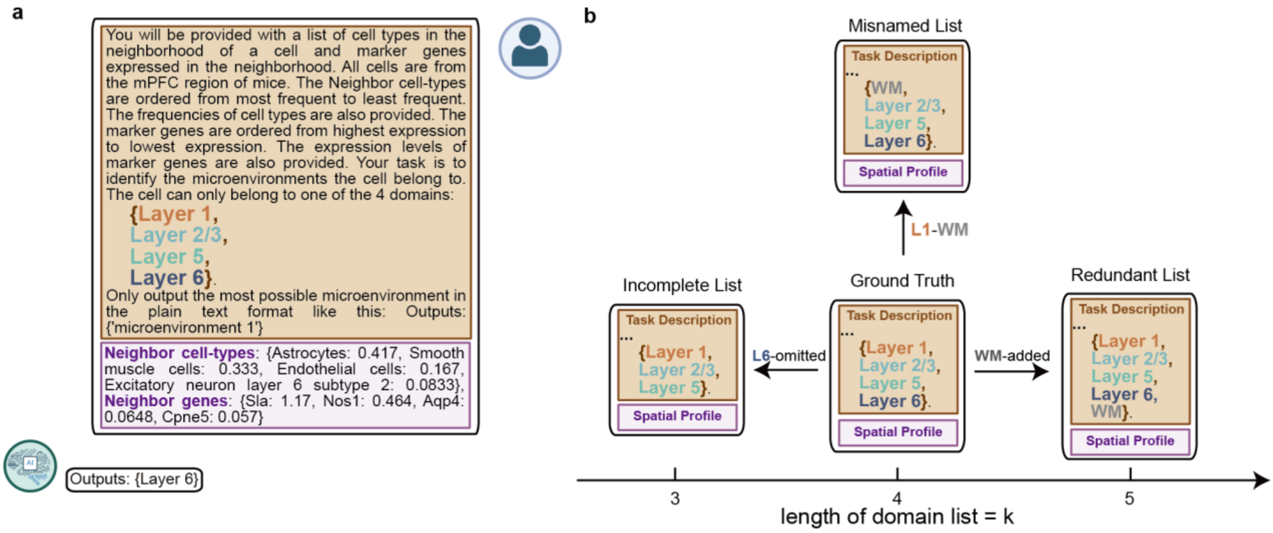


**Supplementary Figure 2.** Domain-list misspecification settings used to evaluate SPEAK. a) Example SPEAK prompt containing the potential domain list for the STARmap dataset, neighborhood cell-type frequencies and neighborhood marker genes. **b**) Schematic of three domain-list misspecification settings: incomplete-list, redundant-list and misnamed-list. In the incomplete-list setting, one ground-truth domain was omitted from the potential domain list, providing only three candidate domains. In the redundant-list setting, the White Matter domain, a biologically meaningful domain that did not exist in the SRT data, was added to the potential domain list, providing five candidate domains. In the misnamed-list setting, one ground-truth domain was replaced with the White Matter domain that was absent in the data.

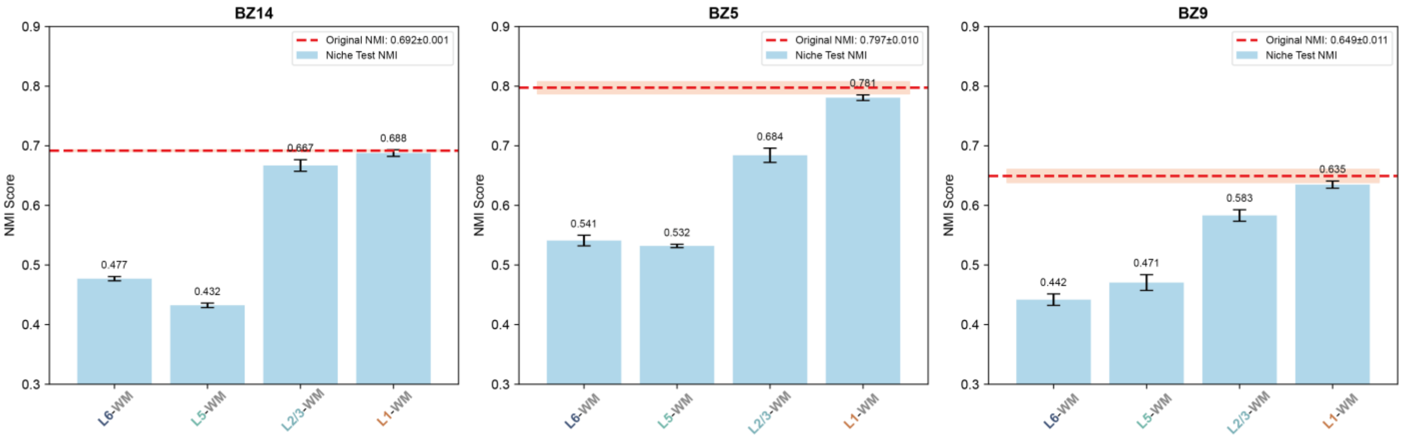


**Supplementary Figure 3.** SPEAK performance under misnamed-list setting for the domain-list misspecification in the STARmap dataset. NMI scores for BZ14, BZ5 and BZ9, when one ground-truth cortical domain was replaced by White Matter in the candidate domain list, were shown in bar plots, in which dashed red lines represent the original NMI obtained with the correctly specified ground-truth domain list and error bars indicate standard error.

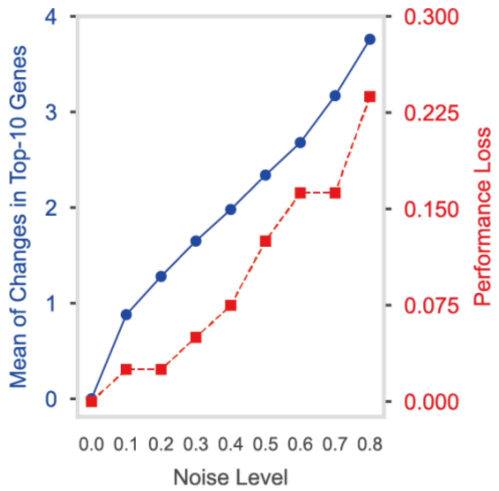


**Supplementary Figure 4.** Robustness of SPEAK to gene-expression drop-outs in MERFISH dataset. The gene-expression profile of a tissue section from MERFISH dataset was perturbed by randomly setting 10-80% of expression values to zero. The blue curve shows the mean number of non-overlapping genes between the top-10 neighborhood marker-gene lists before and after introducing drop-outs. The red curve shows relative NMI loss, defined as 1 – (NMI after dropouts/NMI before drop-outs.

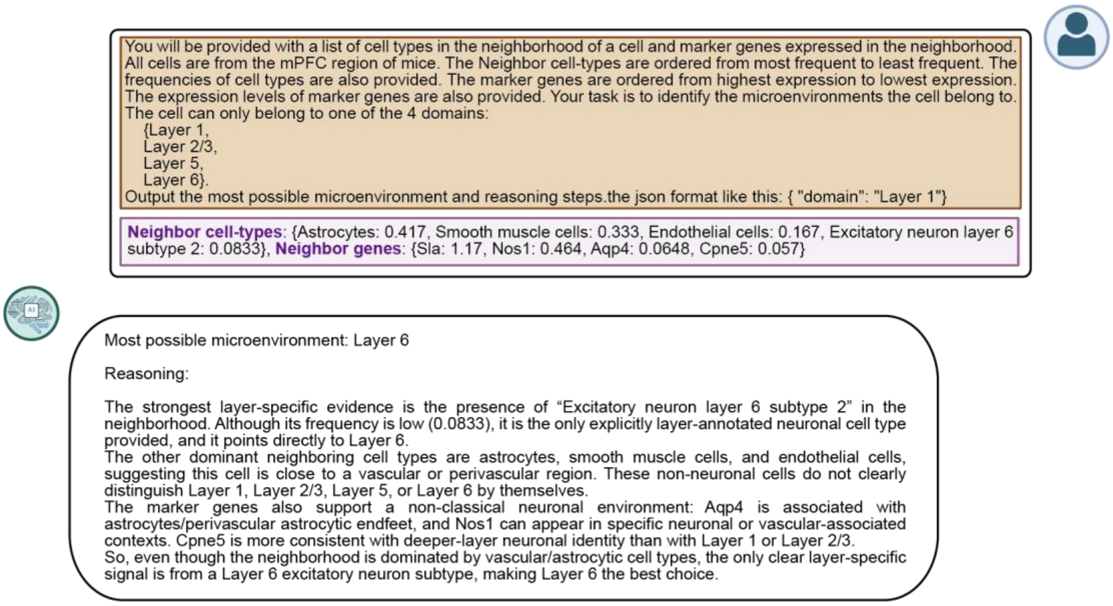


**Supplementary Figure 5.** Example reasoning output from SPEAK using STARMap dataset. Example prompt (top panel) and response showing SPEAK output (bottom panel) with both the predicted microenvironment and the reasoning steps used for the prediction are shown. The prompt includes the candidate cortical domain list, neighborhood cell-type frequencies and neighborhood marker genes. The response assigns the example cell to Layer 6 and provides the evidence used for that assignment.

**Supplementary Table 1.** Normalized Mutual Information (NMI) of different methods in between-section validation on the three benchmarking datasets. Methods were ranked within each dataset and the average rank across datasets was reported. Note that SPEAK-F (within-section) reports the results on cell/spots from the same tissue section that generated the fine-tuning data but not included in the fine-tuning data. It was not included in rank of performance calculation. NMI scores were shown in Mean ± Standard Deviation.

| Method Type | Method | STARmap | Visium | MERFISH | Avg. | Rank |
|---|---|---|---|---|---|---|
| LLM-based | SPEAK-F (within-section) | 0.811±0.003 | 0.759±0.002 | 0.795±0.006 | 0.789 | nan |
| | SPEAK-Fs | 0.752±0.013 | 0.695±0.036 | 0.610±0.027 | 0.686 | 1.7 |
| | SPEAK-F | 0.753±0.011 | 0.678±0.051 | 0.581±0.020 | 0.67 | 2.0 |
| | SPEAK-Z | 0.755±0.040 | 0.487±0.103 | 0.068±0.031 | 0.437 | 10.3 |
| Non-LLM based | BASS | 0.693±0.100 | 0.581±0.021 | 0.519±0.053 | 0.598 | 5.0 |
| | SCAN-IT | 0.630±0.055 | 0.546±0.047 | 0.578±0.045 | 0.585 | 5.7 |
| | BayesSpace | nan±nan | 0.565±0.087 | nan±nan | 0.565 | 6.0 |
| | stLearn | nan±nan | 0.552±0.014 | nan±nan | 0.552 | 7.0 |
| | GraphST | 0.433±0.061 | 0.592±0.049 | 0.317±0.056 | 0.448 | 7.0 |
| | IRIS | 0.310±0.148 | 0.640±0.016 | 0.331±0.063 | 0.427 | 7.3 |
| | SpaceFlow | 0.606±0.061 | 0.433±0.042 | 0.535±0.077 | 0.525 | 9.3 |
| | CCST | 0.353±0.083 | 0.507±0.022 | 0.468±0.031 | 0.443 | 9.3 |
| | Seurat | 0.748±0.095 | 0.474±0.047 | 0.275±0.022 | 0.499 | 9.7 |
| | SpaGCN | 0.318±0.011 | 0.513±0.047 | 0.214±0.015 | 0.348 | 10.3 |
| | STAGATE | 0.538±0.079 | 0.507±0.042 | 0.204±0.085 | 0.417 | 10.7 |
| | SEDR | 0.113±0.057 | 0.532±0.030 | 0.142±0.045 | 0.263 | 12.0 |
| | conST | 0.067±0.021 | 0.511±0.084 | 0.107±0.012 | 0.228 | 13.3 |
| | SpaGCN(HE) | nan±nan | 0.475±0.043 | nan±nan | 0.475 | 15.0 |
| Non-Spatial | Leiden | 0.066±0.026 | 0.329±0.009 | 0.177±0.004 | 0.191 | 15.3 |
| | Louvain | 0.065±0.021 | 0.336±0.014 | 0.169±0.009 | 0.19 | 15.7 |

**Supplementary Table 2.** Homogeneity score (HOM) of different methods in between-section validation on the three benchmarking datasets. Methods were ranked within each dataset and the average rank across datasets was reported. Note that SPEAK-F (within-section) reports the results on cell/spots from the same tissue section that generated the fine-tuning data but not included in the fine-tuning data. It was not included in rank of performance calculation. HOM scores were shown in Mean ± Standard Deviation.

| Method Type | Method | STARmap | Visium | MERFISH | Avg. | Rank |
|---|---|---|---|---|---|---|
| LLM-based | SPEAK-F (within-section) | 0.814±0.003 | 0.737±0.002 | 0.788±0.006 | 0.78 | nan |
| | SPEAK-Fs | 0.757±0.020 | 0.683±0.036 | 0.613±0.021 | 0.684 | 1.0 |
| | SPEAK-F | 0.755±0.018 | 0.662±0.044 | 0.580±0.017 | 0.666 | 2.3 |
| | SPEAK-Z | 0.753±0.043 | 0.429±0.087 | 0.045±0.025 | 0.409 | 12.0 |
| Non-LLM based | SCAN-IT | 0.716±0.077 | 0.561±0.054 | 0.598±0.048 | 0.625 | 5.0 |
| | BASS | 0.715±0.090 | 0.585±0.020 | 0.432±0.043 | 0.577 | 6.0 |
| | BayesSpace | nan±nan | 0.565±0.079 | nan±nan | 0.565 | 6.0 |
| | SpaceFlow | 0.753±0.066 | 0.529±0.048 | 0.483±0.080 | 0.588 | 6.3 |
| | GraphST | 0.440±0.045 | 0.591±0.054 | 0.219±0.042 | 0.416 | 7.3 |
| | IRIS | 0.300±0.162 | 0.627±0.007 | 0.328±0.065 | 0.418 | 7.3 |
| | CCST | 0.360±0.091 | 0.534±0.028 | 0.536±0.035 | 0.477 | 7.7 |
| | stLearn | nan±nan | 0.543±0.009 | nan±nan | 0.543 | 8.0 |
| | Seurat | 0.744±0.095 | 0.480±0.045 | 0.260±0.015 | 0.495 | 9.3 |
| | SpaGCN | 0.344±0.016 | 0.520±0.045 | 0.210±0.013 | 0.358 | 10.7 |
| | STAGATE | 0.564±0.069 | 0.509±0.037 | 0.207±0.087 | 0.426 | 10.7 |
| | conST | 0.072±0.022 | 0.515±0.087 | 0.110±0.012 | 0.232 | 13.7 |
| | SpaGCN(HE) | nan±nan | 0.483±0.043 | nan±nan | 0.483 | 14.0 |
| | SEDR | 0.114±0.058 | 0.455±0.067 | 0.116±0.036 | 0.228 | 14.3 |
| Non-Spatial | Leiden | 0.065±0.026 | 0.334±0.008 | 0.167±0.007 | 0.189 | 15.3 |
| | Louvain | 0.055±0.017 | 0.335±0.014 | 0.157±0.007 | 0.182 | 15.7 |

**Supplementary Table 3.** Completeness score (COM) of different methods in between-section validation on the three benchmarking datasets. Methods were ranked within each dataset and the average rank across datasets was reported. Note that SPEAK-F (within-section) reports the results on cell/spots from the same tissue section that generated the fine-tuning data but not included in the fine-tuning data. It was not included in rank of performance calculation. COM scores were shown in Mean ± Standard Deviation**.**

| Method Type | Method | STARmap | Visium | MERFISH | Avg. | Rank |
|---|---|---|---|---|---|---|
| LLM-based | SPEAK-F (within-section) | 0.808±0.004 | 0.784±0.004 | 0.802±0.006 | 0.798 | nan |
| | SPEAK-Fs | 0.748±0.006 | 0.707±0.040 | 0.608±0.033 | 0.688 | 2.3 |
| | SPEAK-F | 0.750±0.006 | 0.695±0.060 | 0.582±0.025 | 0.676 | 3.3 |
| | SPEAK-Z | 0.757±0.038 | 0.565±0.132 | 0.153±0.036 | 0.492 | 8.0 |
| Non-LLM based | BASS | 0.672±0.107 | 0.577±0.022 | 0.651±0.070 | 0.633 | 4.0 |
| | GraphST | 0.429±0.083 | 0.594±0.044 | 0.593±0.129 | 0.538 | 6.0 |
| | BayesSpace | nan±nan | 0.566±0.096 | nan±nan | 0.566 | 7.0 |
| | SCAN-IT | 0.566±0.057 | 0.533±0.045 | 0.560±0.045 | 0.553 | 7.3 |
| | IRIS | 0.325±0.128 | 0.653±0.025 | 0.335±0.062 | 0.438 | 7.7 |
| | Seurat | 0.751±0.096 | 0.469±0.048 | 0.291±0.032 | 0.504 | 8.7 |
| | stLearn | nan±nan | 0.562±0.022 | nan±nan | 0.562 | 9.0 |
| | SpaceFlow | 0.508±0.059 | 0.367±0.037 | 0.603±0.077 | 0.493 | 9.3 |
| | SEDR | 0.113±0.057 | 0.659±0.047 | 0.187±0.063 | 0.32 | 9.7 |
| | CCST | 0.350±0.082 | 0.483±0.021 | 0.415±0.030 | 0.416 | 10.3 |
| | STAGATE | 0.516±0.087 | 0.506±0.047 | 0.201±0.083 | 0.408 | 10.3 |
| | SpaGCN | 0.297±0.013 | 0.506±0.050 | 0.218±0.018 | 0.34 | 11.3 |
| | conST | 0.063±0.019 | 0.507±0.081 | 0.104±0.012 | 0.224 | 14.3 |
| | SpaGCN(HE) | nan±nan | 0.468±0.042 | nan±nan | 0.468 | 16.0 |
| Non-Spatial | Leiden | 0.067±0.027 | 0.325±0.011 | 0.190±0.004 | 0.194 | 15.3 |
| | Louvain | 0.079±0.029 | 0.337±0.016 | 0.183±0.011 | 0.2 | 15.3 |

**Supplementary Table 4.** Average Silhouette Width (ASW) of different methods in between-section validation on the three benchmarking datasets. Methods were ranked within each dataset and the average rank across datasets was reported. Note that SPEAK-F (within-section) reports the results on cell/spots from the same tissue section that generated the fine-tuning data but not included in the fine-tuning data. It was not included in rank of performance calculation. ASW scores were shown in Mean ± Standard Deviation.

| Method Type | Method | STARmap | Visium | MERFISH | Avg. | Rank |
|---|---|---|---|---|---|---|
| LLM-based | SPEAK-F (within-section) | 0.267±0.000 | 0.018±0.002 | -0.055±0.001 | 0.077 | nan |
| | SPEAK-Fs | 0.226±0.003 | 0.010±0.034 | -0.117±0.034 | 0.04 | 7.0 |
| | SPEAK-F | 0.226±0.005 | 0.001±0.046 | -0.147±0.044 | 0.026 | 9.7 |
| | SPEAK-Z | 0.231±0.037 | -0.065±0.096 | -0.225±0.096 | -0.019 | 11.7 |
| Non-LLM based | BASS | 0.187±0.074 | 0.087±0.022 | -0.017±0.020 | 0.086 | 3.0 |
| | SCAN-IT | 0.184±0.032 | 0.162±0.080 | -0.018±0.056 | 0.109 | 3.0 |
| | CCST | 0.064±0.052 | 0.170±0.078 | 0.292±0.018 | 0.175 | 3.3 |
| | BayesSpace | nan±nan | 0.085±0.070 | nan±nan | 0.085 | 4.0 |
| | IRIS | 0.133±0.116 | 0.045±0.011 | -0.056±0.079 | 0.041 | 6.3 |
| | SEDR | -0.024±0.035 | 0.105±0.070 | -0.121±0.067 | -0.013 | 7.7 |
| | stLearn | nan±nan | 0.044±0.010 | nan±nan | 0.044 | 8.0 |
| | GraphST | 0.054±0.052 | 0.040±0.012 | -0.126±0.035 | -0.011 | 9.7 |
| | STAGATE | 0.111±0.030 | 0.060±0.023 | -0.195±0.044 | -0.008 | 9.7 |
| | SpaGCN(HE) | nan±nan | 0.021±0.035 | nan±nan | 0.021 | 10.0 |
| | SpaGCN | 0.025±0.023 | 0.082±0.043 | -0.172±0.011 | -0.022 | 10.3 |
| | SpaceFlow | 0.080±0.036 | -0.066±0.029 | -0.029±0.059 | -0.005 | 10.3 |
| | Seurat | 0.212±0.052 | -0.001±0.008 | -0.166±0.017 | 0.015 | 10.7 |
| | conST | -0.056±0.009 | -0.001±0.024 | -0.101±0.015 | -0.052 | 12.3 |
| Non-Spatial | Louvain | -0.091±0.010 | 0.010±0.019 | -0.143±0.011 | -0.075 | 12.7 |
| | Leiden | -0.100±0.010 | 0.001±0.010 | -0.168±0.025 | -0.089 | 14.3 |

**Supplementary Table 5.** Spatial chaos score (CHAOS) of different methods in between-section validation on the three benchmarking datasets. Methods were ranked within each dataset and the average rank across datasets was reported. Note that SPEAK-F (within-section) reports the results on cell/spots from the same tissue section that generated the fine-tuning data but not included in the fine-tuning data. It was not included in rank of performance calculation. CHAOS scores were shown in Mean ± Standard Deviation.

| Method Type | Method | STARmap | Visium | MERFISH | Avg. | Rank |
|---|---|---|---|---|---|---|
| LLM-based | SPEAK-Fs | 0.073±0.001 | 0.062±0.002 | 0.030±0.001 | 0.055 | 4.3 |
| | SPEAK-Z | 0.072±0.001 | 0.062±0.002 | 0.030±0.001 | 0.054 | 4.3 |
| | SPEAK-F | 0.074±0.001 | 0.062±0.001 | 0.030±0.001 | 0.055 | 6.3 |
| | SPEAK-F (within-section) | 0.077±0.000 | 0.062±0.001 | 0.034±0.000 | 0.057 | nan |
| Non-LLM based | IRIS | 0.071±0.002 | 0.061±0.001 | 0.029±0.001 | 0.054 | 1.0 |
| | SCAN-IT | 0.073±0.003 | 0.061±0.001 | 0.029±0.000 | 0.054 | 2.3 |
| | CCST | 0.077±0.003 | 0.061±0.001 | 0.028±0.000 | 0.055 | 2.7 |
| | BASS | 0.074±0.002 | 0.061±0.001 | 0.029±0.001 | 0.055 | 4.0 |
| | Seurat | 0.072±0.003 | 0.062±0.002 | 0.031±0.001 | 0.056 | 7.0 |
| | SpaceFlow | 0.074±0.002 | 0.064±0.002 | 0.029±0.001 | 0.056 | 7.7 |
| | GraphST | 0.077±0.002 | 0.063±0.002 | 0.030±0.001 | 0.056 | 9.3 |
| | STAGATE | 0.077±0.003 | 0.062±0.001 | 0.048±0.004 | 0.062 | 10.3 |
| | SEDR | 0.107±0.001 | 0.062±0.001 | 0.046±0.004 | 0.072 | 11.3 |
| | BayesSpace | nan±nan | 0.063±0.002 | nan±nan | 0.063 | 12.0 |
| | stLearn | nan±nan | 0.064±0.001 | nan±nan | 0.064 | 13.0 |
| | SpaGCN | 0.091±0.002 | 0.065±0.002 | 0.049±0.001 | 0.068 | 14.0 |
| | conST | 0.110±0.002 | 0.065±0.003 | 0.060±0.002 | 0.078 | 15.7 |
| | SpaGCN(HE) | nan±nan | 0.067±0.002 | nan±nan | 0.067 | 17.0 |
| Non-Spatial | Louvain | 0.091±0.002 | 0.068±0.003 | 0.055±0.001 | 0.071 | 14.7 |
| | Leiden | 0.101±0.002 | 0.069±0.002 | 0.055±0.002 | 0.075 | 16.0 |

**Supplementary Table 6.** Percentage of Allowed outliers for Segmentation (PAS) of different methods in between-section validation on the three benchmarking datasets. Methods were ranked within each dataset and the average rank across datasets was reported. Note that SPEAK-F (within-section) reports the results on cell/spots from the same tissue section that generated the fine-tuning data but not included in the fine-tuning data. It was not included in rank of performance calculation. PAS was shown in Mean ± Standard Deviation.

| Method Type | Method | STARmap | Visium | MERFISH | Avg. | Rank |
|---|---|---|---|---|---|---|
| LLM-based | SPEAK-F (within-section) | 0.009±0.001 | 0.057±0.006 | 0.027±0.001 | 0.031 | nan |
| | SPEAK-Fs | 0.014±0.001 | 0.048±0.011 | 0.041±0.008 | 0.034 | 4.7 |
| | SPEAK-Z | 0.012±0.004 | 0.064±0.037 | 0.068±0.039 | 0.048 | 6.7 |
| | SPEAK-F | 0.022±0.006 | 0.062±0.017 | 0.053±0.009 | 0.046 | 6.7 |
| Non-LLM based | IRIS | 0.008±0.003 | 0.023±0.001 | 0.025±0.003 | 0.018 | 1.0 |
| | SCAN-IT | 0.025±0.006 | 0.015±0.003 | 0.027±0.003 | 0.022 | 3.3 |
| | CCST | 0.111±0.036 | 0.011±0.003 | 0.005±0.001 | 0.042 | 3.3 |
| | BASS | 0.055±0.020 | 0.029±0.000 | 0.026±0.006 | 0.037 | 4.3 |
| | BayesSpace | nan±nan | 0.053±0.013 | nan±nan | 0.053 | 6.0 |
| | SpaceFlow | 0.050±0.006 | 0.199±0.056 | 0.028±0.005 | 0.092 | 9.0 |
| | Seurat | 0.017±0.004 | 0.138±0.024 | 0.150±0.018 | 0.102 | 9.3 |
| | SEDR | 0.462±0.112 | 0.038±0.011 | 0.392±0.089 | 0.298 | 9.7 |
| | GraphST | 0.158±0.030 | 0.118±0.014 | 0.064±0.028 | 0.113 | 10.0 |
| | STAGATE | 0.089±0.025 | 0.084±0.029 | 0.589±0.100 | 0.254 | 11.0 |
| | stLearn | nan±nan | 0.126±0.012 | nan±nan | 0.126 | 12.0 |
| | SpaGCN | 0.356±0.048 | 0.133±0.028 | 0.590±0.039 | 0.36 | 13.7 |
| | conST | 0.700±0.065 | 0.202±0.148 | 0.847±0.023 | 0.583 | 16.0 |
| | SpaGCN(HE) | nan±nan | 0.228±0.053 | nan±nan | 0.228 | 17.0 |
| Non-Spatial | Louvain | 0.316±0.023 | 0.392±0.074 | 0.568±0.042 | 0.426 | 14.0 |
| | Leiden | 0.552±0.123 | 0.442±0.041 | 0.579±0.036 | 0.524 | 15.7 |
